# A new normalization for the Nanostring nCounter gene expression assay

**DOI:** 10.1101/374173

**Authors:** Ramyar Molania, Johann A. Gagnon-Bartsch, Alexander Dobrovic, Terence P Speed

## Abstract

The Nanostring nCounter gene expression assay uses molecular barcodes and single molecule imaging to detect and count hundreds of unique transcripts in a single reaction. These counts need to be normalized to adjust for the amount of sample, variations in assay efficiency, and other factors. Most users adopt the normalization approach described in the nSolver analysis software, which involves background correction based on the observed values of negative control probes, a within-sample normalization using the observed values of positive control probes and normalization across samples using reference (housekeeping) genes. Here we present a new normalization method, Removing Unwanted Variation-III (RUV-III), which makes vital use of technical replicates and suitable control genes. We also propose an approach using pseudo-replicates when technical replicates are not available. The effectiveness of RUV-III is illustrated on four different data sets. We also offer suggestions on the design and analysis of studies involving this technology.

## Introduction

The Nanostring nCounter gene expression platform has become widely used for research and clinical applications due to its ability to directly measure a broad range of mRNA expression levels without cDNA synthesis and amplification steps [1–4]. However, the “accuracy and reliability of gene expression results are dependent upon the proper normalization of the data against internal reference genes” [5]. Most current normalization methods use spiked-in negative and positive controls and reference (housekeeping) genes, each of which has a good record of effectiveness in related technologies. At present the Nanostring analysis software nSolver and the R package NanoStringNorm [6] provide 84 options for normalizing Nanostring gene expression data: 4 ways of using the negative control data × 3 ways of using the positive control data × 7 ways of using the data on their reference genes.

There will always be random errors and possibly also systematic errors in any measurement process. Systematic errors are also called bias, and arise when the measurement process tends to measure something other than what was intended, e.g. [7]. In many measurement problems, there is a bias-variance trade-off. A correction may well remove bias, but it will also increase the variance of the random errors in the data. An averaging may well reduce the variance of the random errors in the data, but it will also add bias. Most normalizations try to reduce both bias and variance, and will achieve these goals to a greater or lesser extent, arriving at some point in the trade-off. However, the question should not be “Is one normalization generally good, better or worse than another?” Rather, we should ask “Are our goals for this data set better achieved using this normalization rather than another?” In abstract terms, we should ask “Do we come out ahead in the trade-off this time, and can we do better?” Similar considerations apply to the details of any normalization. There is always background, but does subtracting background help in this case? There are no perfect housekeeping genes, but does using a particular set of genes as housekeeping genes help in this case?

This does not mean that we cannot find generally good normalization methods, but that we should focus more on assessing the effectiveness of any normalization with each data set and goal. How do we do this? In this paper not only do we present our novel normalization for nCounter gene expression data, but in discussing its possible value, we also illustrate a number of methods we know for assessing normalizations. They range from statistical summaries such as principal component (PCA) plots [8], relative log expression (RLE) plots [9] and technical replicate agreement (TRA) plots for comparing technical replicates, through to recapitulating known biology.

The normalization method presented here, RUV-III, is a novel extension of previously reported methods, which exploit positive and negative control genes and technical replicates [10, 11]. Here our usage of “negative” and “positive” clashes with that of nSolver. We define a negative control gene as one that is not expected to be affected by the biology of interest in a study, not one that is not expected to be expressed. Similarly, a gene is a positive control gene if its biological response is known, e.g. that it should be differentially expressed, not just that it should be expressed. Although we can and will make use of the spiked-in controls in the nCounter expression assay (which we will call NEG and POS in line with nSolver usage), for us negative and positive control genes should be endogenous, that is, part of the gene expression panel being measured. At times we will regard all genes as “negative controls” and get helpful results, so it is important to reiterate a point made in the preceding paragraph: for us the question will not be “Can all genes be suitable negative controls?” but “Does using all genes as negative controls help?” We will see examples where doing so does help, and others where it does not. Selecting suitable negative controls is the major challenge in using RUV-III.

## RESULTS

### Data sets

To evaluate the performance of the RUV-III method on Nanostring gene expression data, we examined 4 studies comprising one in-house and three different published data sets that had technical replicate samples. The details of each study are given separately below.

### Example 1: Lung cancer study

Our in-house Nanostring data set was part of a study of the expression of DNA repair genes in lung adenocarcinoma (LUAD). The data was generated using 15 nCounter cartridges, with the majority of the normal samples being run in a 15^th^ cartridge a year after the rest of the experiment (Supplementary Fig. 1a). For more details on this study, we refer to the supplementary file and Supplementary Figure 1.

We began our analysis with an examination of the Average Plot (Online Methods) of the raw nCounter data (Supplementary Fig. 2). The spiked-in controls were roughly constant across the cartridges, though occasionally showing substantial variability. The average counts of the housekeeping (HK) genes and the library sizes show a marked decline in the 15th cartridge while the averages of the POS and NEG spiked-in controls are fairly stable across all cartridges. This inconsistency poses a challenge for the nCounter normalization.

In Figure 1a, we present three RLE plots (Online Methods), the first for the unnormalized data. For the nCounter normalized data, we applied a standard option: mean+2sd of the NEG transcripts for background correction, and geometric means for both the POS and the HK transcripts. Lastly, the data was normalized by RUV-III using 500 genes with lowest variance as the negative control set, all technical replicates samples (Supplementary Fig. 1b) and k = 6 (Online Methods). In the RLE plot of the raw data (Fig. 1a) we see substantial differences within and across cartridges, with samples from cartridge 15 standing out. This remains the case with nCounter normalized data, whereas the RUV-III normalization yields much more uniform RLE plots, centred around zero (Fig 1a).

**Figure 1.**
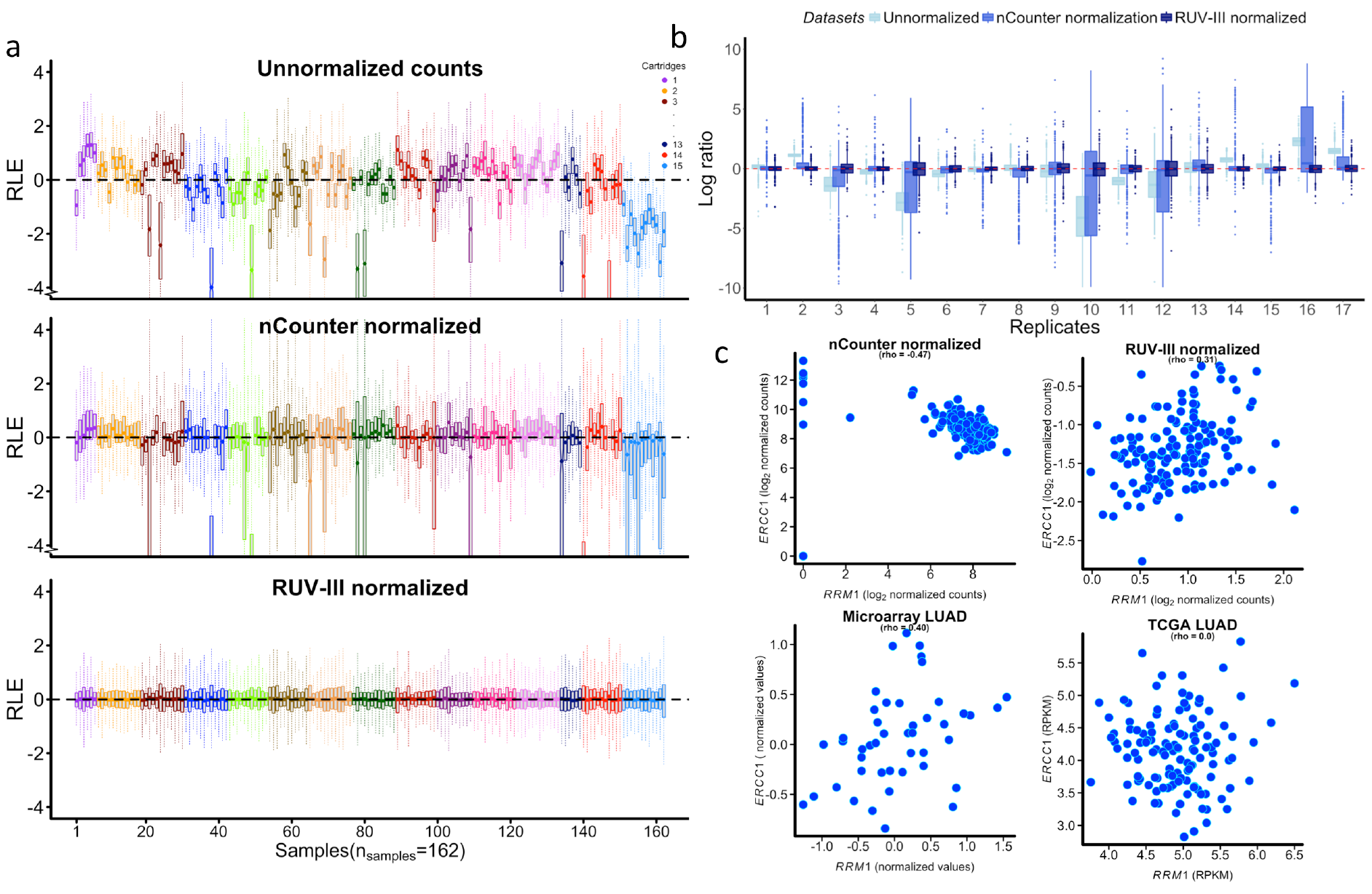
Comparing the performance of nCounter and RUV-III normalization methods. **a)** The relative log expression (RLE) boxplots of unnormalized, nCounter normalized and RUV-III normalized data sets. Ideal RLE distributions would be centered around zero and similar to each other. The boxplots of unnormalized data clearly show substantial variation within and between batches. **b)** Log ratios of genes of all technical replicate samples in unnormalized, nCounter normalized and RUV-III normalized data sets. RUV-III normalization led to smaller log ratios change of genes between technical replicate samples. **c)** Scatter plots of *ERCC1* and *RRM1* gene expression obtained from nCounter normalized data (r =-0.47) and RUV-III normalized data (r = 0.31) and two publically available data sets including Hou *et al*’s [17] microarray gene expression data (r = 0.40) and the TCGA lung adenocarcinoma RNA-SeqV2 (r = 0.0). The nCounter normalization shows very different pattern compared to that of the RUV-III normalized data and the publically available data sets.

We evaluated the similarity of technical replicate samples by producing Technical Replicate Agreement (TRA) plots (Online Methods and Fig. 1b). These show that RUV-III normalization leads to a considerable reduction in the differences between technical replicate samples.

To assess the effectiveness of the different normalizations, we attempted to recapitulate some established biology in our data. A known biological signal in this context is that the expression of the genes *RRM1* and *ERCC1* should be moderately positively correlated [12–15]. In Figure 1c, we present four scatter plots of the expression levels of these two genes. The nCounter normalized data scatter plot clearly deviates from the other three, including the RUV-III normalized data, two of which recapitulate the finding of [12].

We also examined all 84 different nCounter normalization options, displaying in Supplementary Figure 3 the medians of the RLE plots for each sample, the TRA plot and the correlation between *RRM1* and *ERCC1* for all 84 options, and for RUV-III.

Repeating all 84 normalization options on the lung data set using a new set of reference genes obtained by the geNorm [16] method led to results similar to those obtained using the initial set of reference genes (data not shown). In all the assessments we have presented, the RUV-III normalization markedly improves upon the nCounter normalizations.

### Example 2: Inflammatory bowel disease (IBD) study

The study by Peloquin *et al*. [18] of Crohn’s disease (CD) and ulcerative colitis (UC) included samples from 3 tissues (colon, rectum, and terminal ileum) under each of 5 different conditions: CD-uninflamed, CD-inflamed, UC-uninflamed, UC-inflamed and healthy. The cartridges were run in three batches (CodeSets) numbered 2, 3 and 4, see (Supplementary Fig. 4a) and the supplementary text for fuller details.

First we examined the Average and RLE plots of the unnormalized data (Supplementary Fig. 5a, b). Both show noticeable batch differences. The RLE plots of the data normalized by Peloquin *et al* show that there is still noticeable variation within and between batches (Supplementary Fig, 5.a), and this is strongly supported by the batch-colored PCA plot (Fig. 2a). There one of the two clusters is dominated by samples from batch 3, consistent with a batch difference rather than the tissue difference we would expect to see biologically. Interestingly, samples from batch 3 do not stand out if we make our PC2 against PC1 plot with the correlation matrix [19]. However, batch 3 does stand out in PC3 against PC1 plot using the correlation matrix (Supplementary Fig. 6).

**Figure 2.**
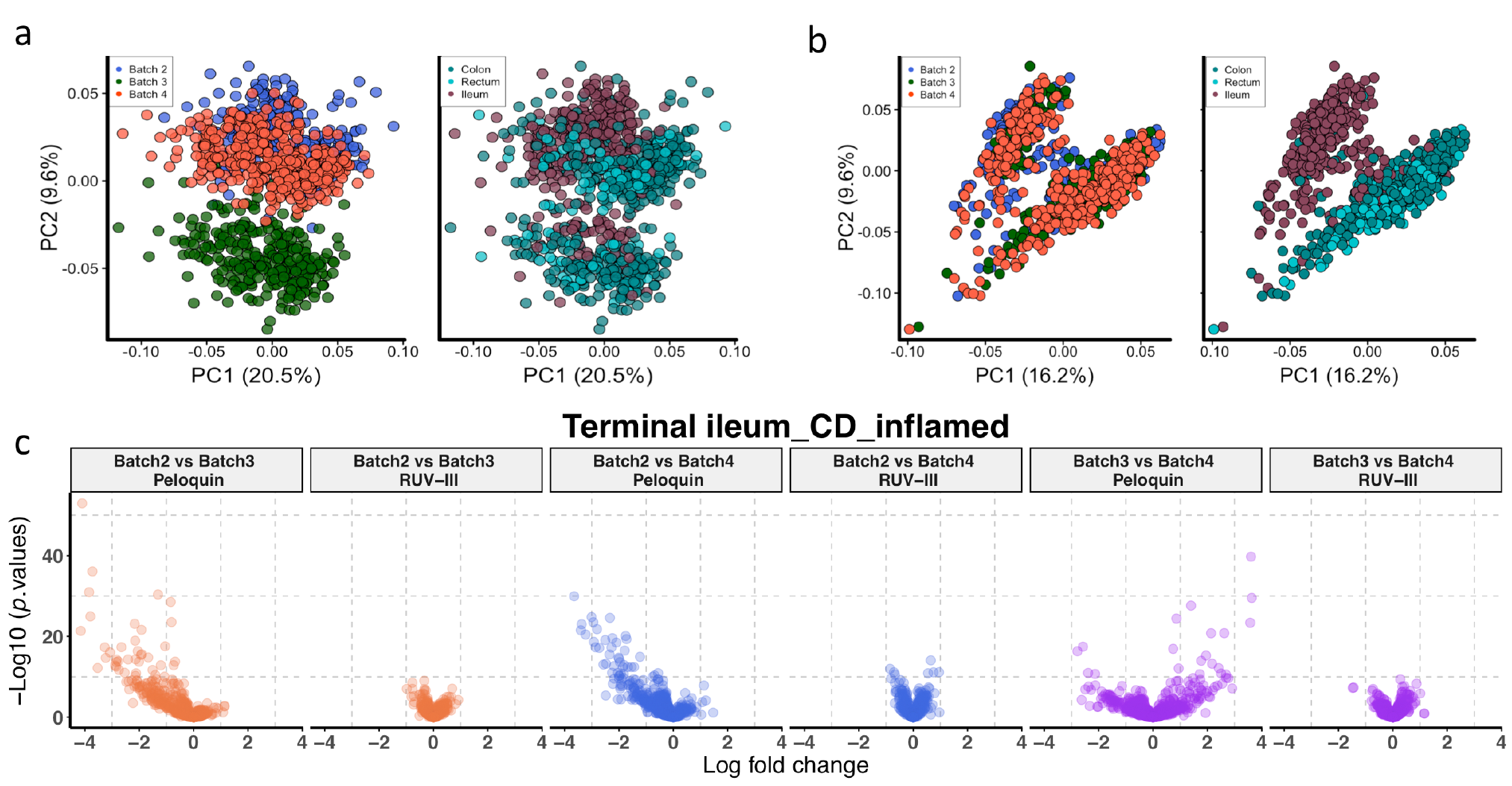
Comparing the performance of different normalization methods on Nanostring data for the inflammatory bowel disease study. **a)** Scatter plots of first two principal components (log scale and centered) for Peloquin normalized Nanostring data colored by reagent lots (left) and by different tissues (right). The second principal component captures all samples of batch 3, which clearly demonstrates that batch effects remain. **b)** Same as **a**, for RUV-III normalized Nanostring data colored by reagents lots (left) and by different tissues (right). The different tissues cluster as expected biologically. **c)** Differential expression analysis of terminal ileum individuals with inflamed Crohn’s disease between pairs of Nanostring batches.

We then normalized the data by RUV-III using all genes as negative controls, all technical replicates (Supplementary Fig. 4b) and the maximum value of K (Online Method). The RUV-III normalization led to RLE plots that were more uniform and centered around zero, and also to the separation of ileum from colorectal tissues in PCA plots (Fig. 2b). This is now consistent with what is known about the biology of these tissues [20].

In order to determine whether the batch differences visible in Figure 2a are of any consequence, we performed a differential expression analysis for each disease state between pairs of batches, comparing the data normalized by Peloquin *et al* with that normalized by RUV-III. The results (Fig. 2c) are displayed in the form of volcano plots with -log_10_(*p*-value) plotted vertically against the log(fold-change) horizontally for each gene counted. More such plots can be found in Supplementary Figure 6. Ideally we should see little evidence of differential expression in these plots, whereas we see a lot in the data normalized by Peloquin *et al*, far more than in the RUV-III normalized data.

Noble *et al* [21]reported microarray gene expression data profiling colon tissue in some of the same states studied in Peloquin *et al*. This enabled us to compare the two normalization results using an orthogonal platform.

We note that the batch effects remaining in the data normalized by Peloquin *et al* (Fig. 3a) can influence downstream analyses. For example, these data suggest that the *CCDC101* gene is up-regulated in UC uninflamed colon compared to normal colon tissue, whereas we do not see this in either the RUV-III normalized data or the Noble *et al* microarray data (Fig. 3b). In Supplementary Figure 8 we see similar results for 6 other genes. The residual batch effect in the data normalized by Peloquin *et al* clearly matters, and it is not present in the RUV-III normalized data.

**Figure 3.**
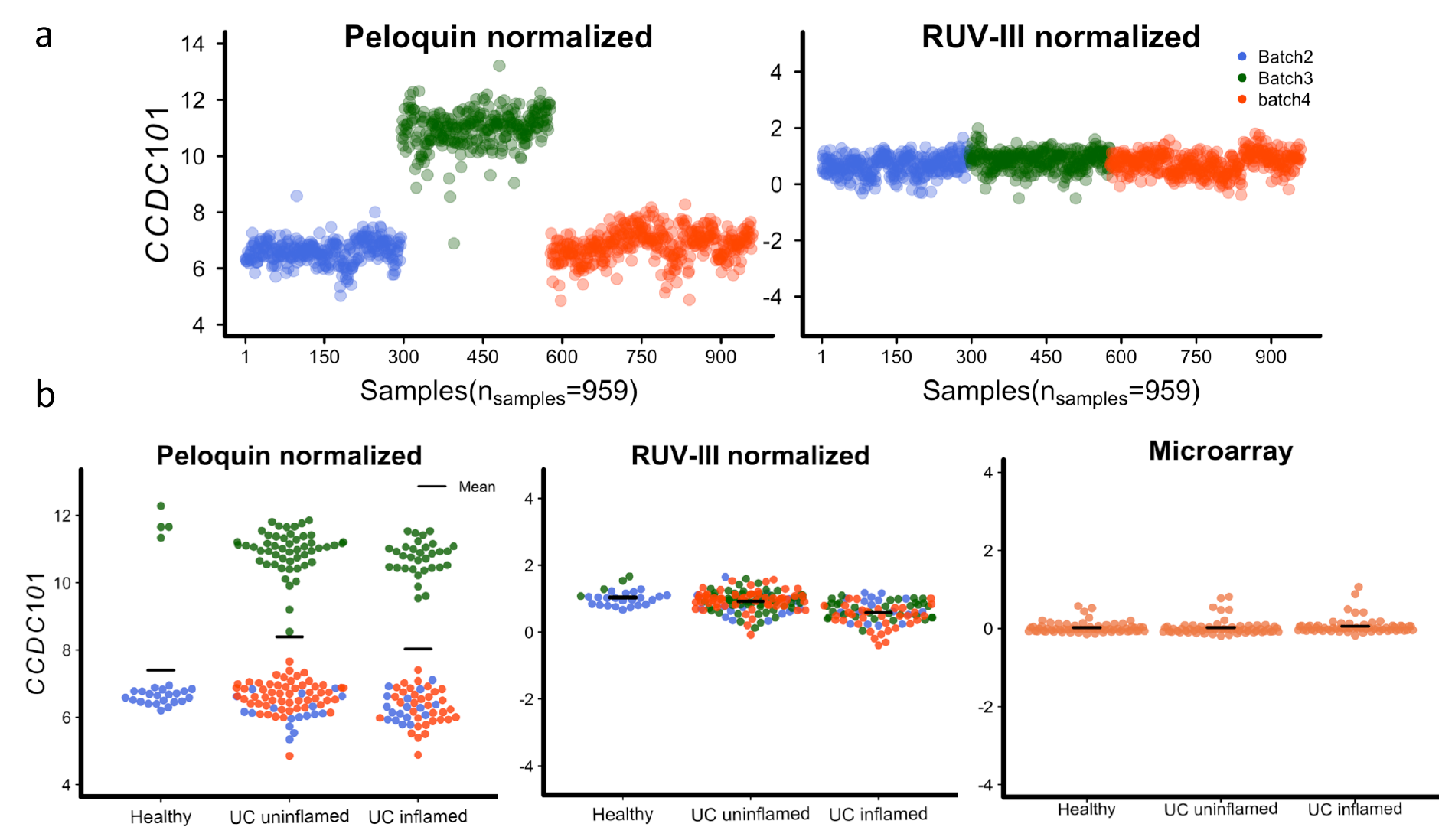
Impact of batch effects on differential expression analysis. **a)** Expression pattern of *CCDC101* gene across different batches of reagents. The Peloquin normalized Nanostring data showed a substantial difference between batch 3 and other batches whereas RUV-III largely removes this unwanted variation. **b)** The expression pattern of the *CCDC101* gene across different the disease states of colon samples. The gene is apparently up-regulated in colon samples in the uninflamed ulcerative colitis state compared to healthy individuals in the Peloquin normalized data, whereas there is no evidence of differentially expression in the RUV-III normalized data or the Noble *et al* microarray data.

### Example 3: Human T cell activation

For a discussion of the study by Ye *et al* [22] of CD4^+^ T cell conditions including activated 4 hours, activated 48 hours, INFβ 4 hours, Th17 48 hours and unstimulated T cells for 4 hours, we refer to the Supplement. The Nanostring data consisted of 1,788 assays generated over 4 months in 2012 and 2013 (Supplementary Fig. 9a). We found only 46 technical duplicates samples, distributed across Nanostring cartridges in 2013 only (Supplementary Fig. 9b).

The Average Plot (Supplementary Fig. 10) of the unnormalized data show a clear downward shift in expression values from 2013 compared to 2012 (hereafter called time effects); this is most clear in the positive spike-in controls. The RLE plots of the same data show that the majority of samples with 48 hour conditions were shifted up compared to those with the 4 hour conditions (Supplementary Fig 11a), and the individual expression patterns of housekeeping (Supplementary Fig. 12a) and gender genes (data not shown) confirm this. The pattern observed in the raw data is strikingly reversed in the data normalized by Ye *et al.* (Supplementary Fig. 11a), where samples under the 48 hours conditions now have generally lower values compared to those with 4 hour conditions, and this too is reflected in individual gene expression patterns (Supplementary Fig. 12b). This is likely the result of using housekeeping genes for normalization across cartridges. We retrieved the authors’ normalized gene expression microarray data from GEO, and produced RLE plots for the 236 genes used in their nCounter assays (Supplementary Fig. 11b). These RLE plots are very different from those of the Nanostring data normalized by Ye *et al.*

Our earlier strategy of using all genes as negative controls and all technical replicates failed to produce a reasonable RLE plot, so we used the 9 (from 236) genes with lowest variance in the microarray data as a set of negative controls and all technical replicates to normalize the Nanostring data using RUV-III (k = 1). The RUV-III normalization largely removed the technical variation between the T cell conditions, but the time effects remained in the data due to not having technical replicate across 2012 and 2013. Enlarging the number of negative control genes from 9 to 100 produced little change (Supplementary Fig. 11a).

Next, we carried out a differential expression analysis between pairs of conditions to examine the concordance between log fold-changes found using the data normalized by Ye *et al* and the RUV-III normalized data and the corresponding results from the microarray data. The result for one of 10 pairwise comparisons is seen in Figure 4a while those for the other 9 are in the supplement (Supplementary Fig. 13). This was repeated for the 100 least variable genes (Supplementary Fig. 14). In 17/20 comparisons, the RUV-III normalized data gives results closer to those from the microarray data.

**Figure 4.**
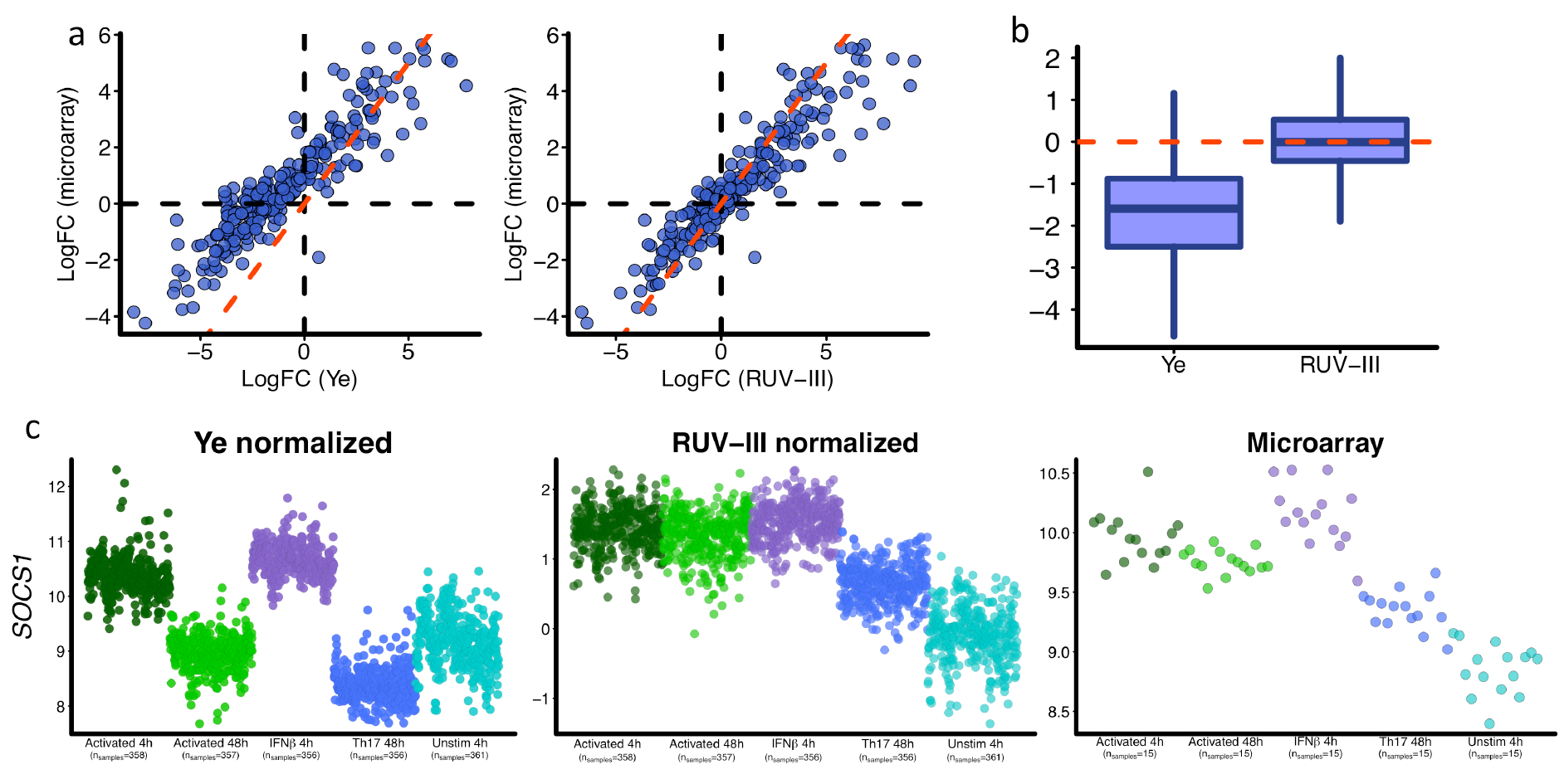
Differential expression analysis between activated 48h and unstimulated 4h conditions and the expression pattern of *SOCS1* gene. **a)** Scatter plots of log fold changes obtained by differential expression analysis between activated 48h and unstimulated conditions for Ye-normalized data and RUV-III normalized data, each compared to the corresponding comparison from Ye *et al* microarray data. The dotted red line is the 45^°^ line. **b)** Boxplot of log fold change (Ye) – log fold change (microarray) and log fold change (RUV-III) – log fold change (microarray). **c)** Expression pattern of *SOCS1* gene across all the T cell conditions in Ye-normalized data, RUV-III normalized data and Ye *et al* microarray data.

We also display several genes whose expression levels differ across the conditions, and compare the patterns of differences seen in Ye-normalized, RUV-III normalized, and the microarray data (Fig. 4b, Supplementary Fig. 15)

In all of these comparisons, the RUV-III normalized data showed better agreement with the microarray data than did the Ye-normalized data.

### Example 4: Dendritic cell study

The final data set we examined is from the study by Lee *et al.* [23] of four different CD14^+^ T cell conditions, unstimulated for 0 hour (UNS 0h), *E. coli* bacterial lipopolysaccharide for 5 hours (LPS 5h), influenza virus for 10 hours (FLU 10h) and interferon beta (INFβ) for 6.5 hours. We refer to the supplement for fuller details, particularly Supplementary Figure 16b for sample collection times and Supplementary Figure 17 for the TRL plot.

We began our analysis with average and RLE plots of the raw and Lee-normalized data (Supplementary Fig. 18a, 19), and we saw that the UNS samples really stood out. Not only were they very spread out in time (Supplementary Fig. 16b), we noticed that they consisted of 165 that were assayed under the UNS condition alone (U), 276 that were assayed under all four conditions (ULVI), with smaller numbers for another four types (Supplementary Fig. 16a). We were led to define 11 sample usage types, see supplement for a full explanation of this term, and Supplementary Figure 16a for their numbers.

In figure 5a we present PCA plots of the Lee-normalized data stratified by condition and colored by gender (row 1) and by usage type (row 2), while Supplementary figure 20 and 20a (middle panel) have the same data with the initially RUV-III normalized data.

**Figure 5.**
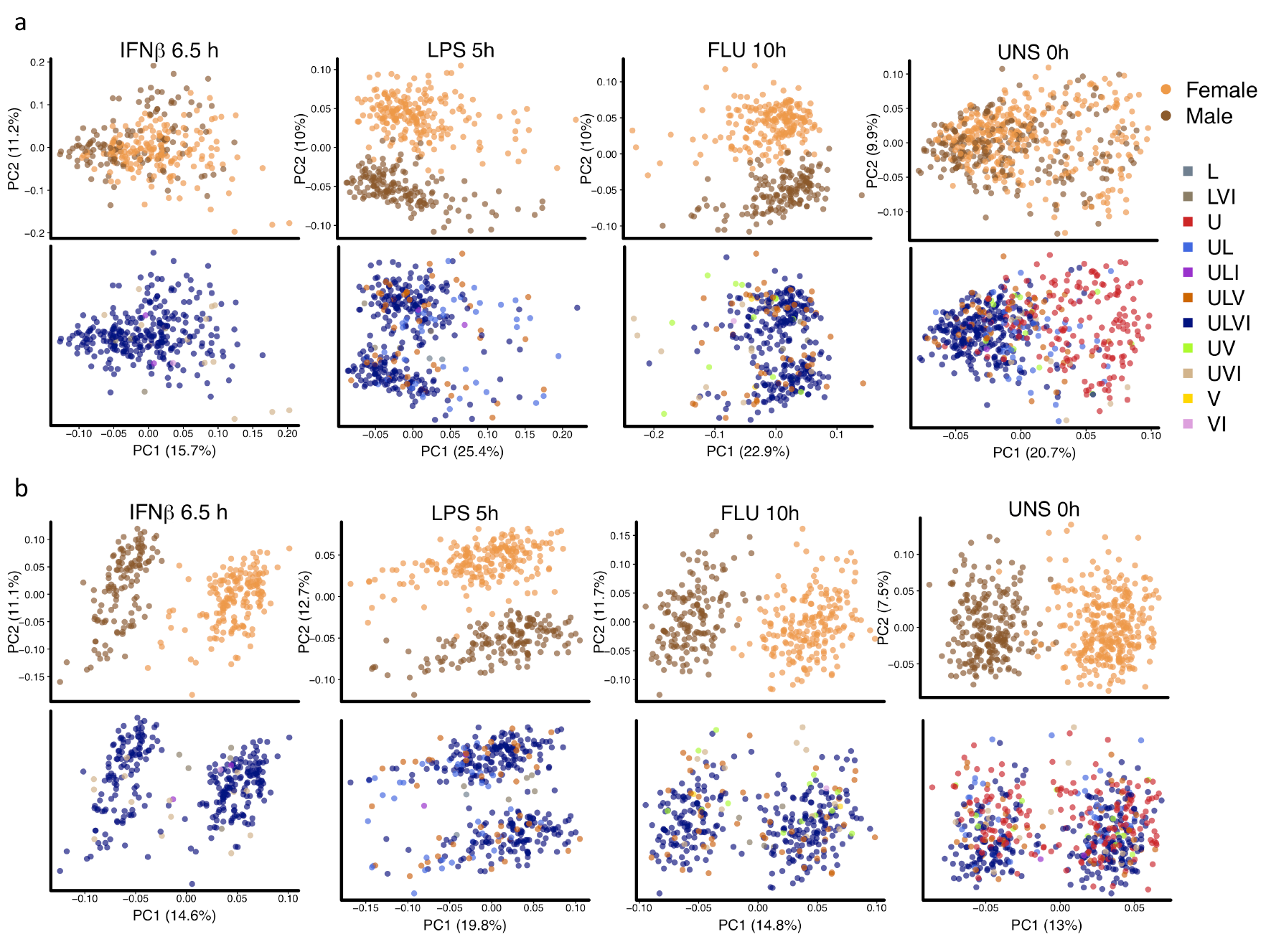
Scatter plots of first two principal components of Lee-normalized data and RUV-III normalized data with all technical replicates and 10 pseudo-replicate pairs. **a)** Principal component analysis of each condition colored by gender (first row) and by usage type (second row). Ideally, PCA plots within condition should have two clear clusters corresponding to males and females, preferably defined by PC1. **b)** Same as **a**, for RUV-III normalized data using all technical replicate samples and 10 pairs of pseudo technical replicate across U and ULVI types of the UNS condition.

A differential expression analysis comparing the U-samples with the rest of the samples showed striking heterogeneity of Lee-normalized gene expression data under the UNS condition (Supplementary Fig. 21a). By definition, there cannot be replicate pairs spanning the U and the ULVI sample usage types, so we created 10 carefully matched pseudo-replicate pairs that did so (Online Methods). We then used RUV-III with all technical replicates and the 10 pseudo-replicate pairs, the 340 least variable genes as negative controls, and k = 10. The resulting RLE plot is in Supplementary figure 19 and the corresponding PCA plots are in rows 3 and 4 of Figure 5.

When we normalized using RUV-III with the pseudo-replicates spanning the U and ULVI sample usage types, the resulting PCA plots split into two clusters on PC1 corresponding to gender for 3 of the 4 conditions (Fig. 5b). It seems likely that there is heterogeneity under the LPS 5h condition which could be removed by creating suitable pseudo-replicates that would lead to a gender split on PC1 for that condition too, but we did not explore that. These clusterings, the RLE plots in Supplementary Figure 19 and the greatly reduced gene expression heterogeneity evident in Supplementary Figure 21c demonstrate the value of using pseudo-replicates with RUV-III.

## DISCUSSION

Normalization is key to deriving meaningful biological results from complex gene expression data sets, especially when the data is compromised by unwanted variation. A normalization method for Nanostring gene expression data must be able to adjust for sample loading differences, variations in assay efficiency, the use of different Nanostring CodeSet batches, and other factors.

Our goal in this paper is to compare our new way of normalizing Nanostring data based upon the use of technical replicates with other normalizations in the literature. In order to do so, we made use of our own data on lung adenocarcinoma samples and three published Nanostring data sets. Our comparisons are based on statistical summaries, biological positive controls, and concordance with previous results.

In the lung cancer study, in the summary of all 84 nCounter normalization options (Supplementary Fig. 3), NNN and some other options have reasonable narrow TRA plots and reasonable (*RRM1, ERCC1*) correlation close to that of the RUV-III normalization, but RLE plots for these options include many boxes far from centered (blue), while with RUV-III they are all centered (Fig. 1a, and extreme right of Supplementary Fig. 3).

Peloquin *et al* [18] compared their results for the healthy patients with the corresponding ones from the microarray data, and found concordance. We note that all but 4 of the normal samples in their comparison were from batch 2 (Supplementary fig 4), and that if they had used other groups in this comparison, the agreement may not have been so good.

We noticed that the different experimental conditions were not uniformly distributed across the 167 Nanostring cartridges (batches) in the T cell study by Ye *et al*. The majority of samples in the 48 hours condition were run together, and the majority of samples with 4 hours conditions were also run together (Supplementary Fig. 9a). This suggests that the study design may have led to a confounding between biological factors of interest and the Nanostring cartridges, though we cannot offer an explanation for what we see. The patterns of expression across conditions seen in the housekeeping genes from the microarray data did not resemble that seen in either the raw or the Ye *et al-*normalized data (Supplementary Fig 12).

Although there were plenty of technical replicates, these cannot bridge conditions, and so we had to rely more heavily on negative control genes for RUV-III. We had great trouble finding suitable ones using just the Nanostring data, but the 9 genes exhibiting least variation across conditions in the microarray data worked well, and replacing 9 by 100 was almost as good.

There are some quite noticeable differences between the log fold changes obtained using the Ye normalization and the microarray data, whereas the agreement between the RUV-III normalized data and the microarray data is much better (Figure 4a, Supplementary Figs. 13-14). We conclude that the Ye normalization has shortcomings, whereas using RUV-III, we obtain much better concordance with the microarray data, and that this normalization greatly reduced, but did not completely remove the impact of the likely confounding effects.

The challenge normalizing the data from the dendritic study by Lee *et al* resembled that just discussed. We felt that a good normalization of all the data should lead to PCA plots clearly separating the four conditions, and that PCA plots within condition should have two clear clusters corresponding to males and females, ideally defined by PC1. The Lee-normalization did not fully achieve this (Fig. 5a), and neither did our initial RUV-III normalization (Supplementary Fig. 20). A major obstacle was the unstimulated (UNS) condition, which turned out to be extremely heterogeneous, consisting of samples with very different collection times and usages (Fig. 5a, Supplementary Fig. 16). The heterogeneity was dramatic enough for there to be many genes extremely significantly differentially expressed between samples in the UNS condition that were only measured under that condition and those what were measured under the other three conditions as well. We can offer no explanation as to why this might be so, but careful choice of pseudo-replicates bridging these two subsets of samples led to Figure 5b, see also Supplementary Figure 21.

We will discuss design issues more fully elsewhere, but certain clear principles should have emerged from the above discussion. The inclusion of carefully chosen endogenous negative control genes in the code set, say 10% of all genes, will more than repay the extra expense. Likewise, the strategic inclusion of at least 10% technical replicates, beginning with at least one in each cartridge, will suffice to permit RUV-III to do its job. Exactly what “strategic” means in any given study will depend on the nature of the inevitable unwanted variation, whether it be CodeSet batch, as with the IBD data, time as in the T cell and dendritic cell data, or other highly specific features such as in the last mentioned, which might be hard to anticipate, but definitely need to be considered.

If an nCounter gene expression data set has very few or no replicates, RUV-III cannot be used. In such cases, analysts might care to use our approach to assessing their own normalization, one of the 84 nCounter options, or one of the methods presented in [11] which do not require replicates.

In summary, we have shown that the use of technical replicates, pseudo-replicates where necessary, and suitably chosen negative control genes in can RUV-III lead to satisfactory normalization of Nanostring data where other methods exhibit shortcomings. We have shown a variety of ways of assessing normalizations, and inter alia, given hints for better designing largish Nanostring studies. As shown in the examples we presented, RUV-III can profitably be used to re-analyse existing data sets to yield new insights into the data.

## ONLINE METHODS

### Average plots

These are obtained by taking the *average log count* of all 8 NEG transcripts, of all 6 POS transcripts, and of all of the housekeeping genes, together with library size in one nCounter assay, and plotting these three averages and the *log(total count)* for that assay, all suitably offset vertically to minimize overlap, in the order in which the assays were run, or related order. Trends, temporal clustering, or other patterns in these plots can reveal issues that normalization should address.

### Technical Replicate Location (TRL) plots

Here we depict the location of these replicates in time, and in relation to known batches, using lines connecting replicate pairs, triples, etc. When connecting lines are viewed together with Average and RLE plots (see below), we can get a sense of how well differences between replicates can help remove batch effects. From seeing where there are no connecting lines, we can also get a sense of how suitably defined pseudo-replicates (see below) might help.

### Relative Log Expression (RLE) plots

A common representation of the results of studies like those discussed in this paper is the collection of boxplots of the log[count]s from each assay, lined up in the order in which the assays were run. Again this permits trends, temporal clustering or other patterns to be visible, and it allows outliers to stand out. The *relative log expression* (RLE) plot is a more sensitive modification of this representation, as it removes the variation between mean gene counts. Instead of making a boxplot of all the log[count]s in each assay, the quantities going into the boxplots are relative to the gene-wise median counts. More fully, instead of log(count), we use log(count) – median[log(count)], where for each transcript (gene), the median over all the assays is taken and its log subtracted. Equivalently, the log[count] for a given transcript, is median corrected, where we use the median rather than the mean to reduce the influence of extremes. For a fuller discussion of RLE plots, see [9].

### Principal Component Analysis (PCA) plots

The principal components (also in this context called singular vectors) of the sample × transcript array of log[count]s are the linear combinations of the transcript measurements having the largest, second largest, third largest,… variation, standardized to be of unit length and orthogonal to the preceding components. Our PCA plots are of the second vs the first principal component (PC) of the sample x transcript array. The calculations are done on mean-corrected transcript log counts, using the R code adopted from EDAseq R package [24].

### Technical Replicate Agreement (TRA) plots

If two assays 1 and 2 are technical replicates, that is, independent assays of two samples of cells from the same tissue sample or subject, then for any transcript (gene) we can form the log ratio log[(count_1_ /count_2_)] of the counts from the two assays. These log ratios can then be pooled across all transcripts in the assay, for a given pair of replicates, and across all technical replicate pairs in a study, and a boxplot created out of them. This we call a *technical replicate agreement* (TRA) boxplot. The boxplot, in particular its inter-quartile range, summarizes the average closeness of technical replicates. It provides a simple visual metric for assessing and comparing normalizations. If, as in RUV-III and some other normalizations, explicit use is made of technical replicates, then the contribution of a given replicate pair to the TRA plot must be calculated after normalization without that pair being used, i.e. with them left undeclared as replicates. All TRA plots calculated after RUV-III in this paper were formed this way.

### Differential expression (DE) analysis

Differential expression analyses of all normalized Nanostring data sets in this paper were performed using limma R package [25]. The design matrix was created using the *model.matrix* function for only biological factor of interests. The empirical Bayes method included in limma package were used to carry out differential expression analysis and the unadjusted *p* values and log fold changes were used for further analysis.

### Volcano plots

A volcano plot is a summary of the results of a two-condition differential expression analysis, where each point corresponds to a transcript, the vertical axis being −*10log_10_p* and the horizontal axis *log(FC)* for that transcript, where *FC* is the fold change and *p* denotes the associated (unadjusted) *p*-value for the test of the null hypothesis of no change. These plots provide a convenient visualization of the efficacy of the study and the location of controls; ideally negative controls should be near (0,0), while positive controls should be well away from 0 on both axes.

### *p-*value histograms

In a two-condition differential expression study with hundreds or thousands of transcripts (genes), the majority will in general not be differentially expressed (DE), in which case their (unadjusted) *p*-values will be uniformly distributed over (0,1), while those transcripts (genes) that are DE will have *p*-values close to zero. As a result, we usually expect that the histogram of *p*-values from a DE study should look uniform, with a possible spike near zero. The presence of unwanted variation will frequently destroy this expected shape. If the number of transcripts (genes) is just a few hundred, and they were pre-selected to exhibit the DE under study, then the expected shape may not result, even in the absence of unwanted variation, and so extra care in interpretation is needed in this case.

### Selection of negative control genes

First, we need to reiterate that in our usage, a negative control gene is one that is *not expected to change much* across the samples being analysed. Second, we need to emphasise that our approach to negative controls is pragmatic: if using a given gene in RUV-III as one of the set of negative control genes helps, as indicated by various measures, then whether or not it is an *ideal* negative control gene is moot: it helped. This does not rule out the fact that we may be able to do better by replacing that gene with a different gene designated a negative control. Thirdly, we point out that theoretical analyses not presented here show that it is not the extent to which individual genes designated as negative controls are ideal or less than ideal negative controls that drives the success or otherwise of RUV-III; that is a property of the full set of negative controls. We will frequently get very good results using the entire set of genes being studied as negative controls, even when many genes are changing. (That this is not unreasonable follow from the theory just mentioned.) Fourthly, it is usually the case that using more genes as negative control genes is better than fewer. That is a matter of stability, but as stated in the introduction, there is a bias-variance trade-off here: using too many genes as negative controls may be counter-productive. You must look and see. Fifthly, endogenous genes generally make more suitable negative controls than spike-ins. The reason here is the obvious one, namely, that endogenous genes have shared the complete sample experience of the other genes, whereas spike-ins can only reflect unwanted variation in the process from the point at which they were added onwards. The best source of endogenous negative control genes are ones that were found to be stable in previous studies similar to the one being analysed. Given that Nanostring expression assays are frequently designed following initial, genome-wide microarray or RNA-seq assays, it should be straightforward to include 20-50 endogenous likely negative controls. We believe that the results in this paper show that their presence for use in normalizing the data will definitely justify the extra expense. Suitable generic sources of negative controls include those genes that are traditionally regarded as housekeeping genes (*ACTB*, *GAPDH*, etc). Gender related genes, such as those on the Y chromosome, can double as negative and positive controls in a gender DE. Having said this, it can at times be quite challenging to find suitable endogenous negative controls in Nanostring studies, especially where there are few “housekeeping” genes, and where no prior studies are available. Sometimes it becomes a matter of trial and error, of doing the best you can.

### RUV-III method

In RUV-III we make use of two separate sources of information on the unwanted variation in a study with technical replicates: that embodied in the differences between the expression values of the replicate samples, and that embodied in the expression values of the negative control transcripts (genes). Informally, the three steps that make up RUV-III are the following:

i) take residuals from the replicate expression measurements and estimate one aspect of the unwanted variation;
ii) take the results of i) together with the expression values of the negative controls, and estimate another aspect of the unwanted variation; and
iii) combine the results from i), ii) into an estimate of the unwanted variation, and subtract that from the data.

The dimension (k) of the unwanted variation also needs to be determined. Full details of the method and R code are given in the Supplement Method and ruv R package (https://cran.r-project.org/package=ruv) respectively.

### Choice of k

To use RUV-III, we need to fix the number k of (linear) dimensions of unwanted variation to be removed. As with the set of negative controls, and the replicates (or pseudo-replicates), there are no clear rules for determining k. Our general approach is to repeat the analysis with different values of, and then evaluate the quality of each analysis using various statistical metrics and prior biological knowledge, as explained in the above. RUV-III is generally robust to overestimating k, but not always. If the number of negative control genes is less than the number of samples, the maximum value of k is equal to number of technical replicates (m-m’). We refer to comments on RUV-III in the Supplementary Methods.

### Number and placement of technical replicates, and the occasional need to define pseudo-replicates

When a study has several batches, it is highly desirable to have replicates placed so that the unwanted variation between any pair of batches can be captured, directly or indirectly, via differences of expression values between technical replicates. For example, if there are 3 batches, and (for instance) there are samples with technical replicates present in all 3, then all gene-wise batch-batch differences can be estimated using differences of these technical replicate samples. (How well is a different matter, see later.) But we need not have samples with technical replicates in all 3 batches: some which span batches 1 and 2, and others which span batches 2 and 3 will suffice, for the difference between batches 1 and 3 can be estimated by the sum of a 1-2 and a 2-3 difference. Again how well it can be estimated is a different matter. This observation generalizes in a natural way, which we do not spell out at the moment. Roughly, the best results will come when different batches can be “connected” by a short chain of replicate pairs. To see this point, suppose that we have two batches, and that all the technical replicates lie wholly within batch 1, or wholly within batch 2, with no replicate pair linking batches 1 and 2. How then can we adjust all gene expression measurements for any bias in the batches? The answer is that we can’t do so directly for all genes, as we could when the two batches were “connected”, we can only get a direct measure of the 1-2 difference for our negative control genes. When the two batches are not connected by a replicate pair, we may do a poor job removing batch differences using RUV-III.

What can be done in this case? Suppose that there is a pair of samples with one in batch 1 and the other in batch 2, and that we believe that the batch differences between these two samples of most genes are greater than the biological differences. We might have reason to believe that the batch 1-2 difference is really large, for example, by looking at the RLE plot. If we designate this pair of samples a replicate pair, and use RUV-III, we may find ourselves coming out ahead. That is, we may be able to use the standard battery of graphical diagnostics and positive controls to convince ourselves that the resulting estimate of the batch 1-2 difference, while imperfect, was near enough to the unwanted variation we sought to remove, to permit a successful application of RUV-III. We would call the pair of samples so used *pseudo-replicates.* This is precisely what we did with the dendritic cell example.

### Not quite anything goes, but almost, as long as we check

In the preceding discussion, we have suggested that there is no such thing as a perfect negative control, but rather, plausible candidates should be tried, and their value assessed. And we said that, up to a point, the more the better. We have also said that to get RUV-III working well, we’d like replicates suitably spanning the major batches or batch-like groups of samples. For example, if there are visible trends in the average or RLE plots, then we’d like replicates spanning the trend, not all at one end, or all in the middle. When there are samples in excess, and clearly major pockets of unwanted variation, then creating pseudo-replicates may help. All of which brings up a point that cannot be emphasised too much: Any normalization should be critically assessed using all the tools that can be brought to the task: RLE, PCA, TRA, volcano plots, p-value histograms, and comparisons with as much prior information as possible, including positive controls. Normalizing using RUV-III with different sets of negative controls, or making use of pseudo-replicates, if the natural controls or replicates available seem inadequate, is a reasonable approach, which can “rescue” data that is otherwise fatally compromised by unwanted variation.

## Software availability

RUV-III is implemented in the open-source R package ruv, with source code freely available through the CRAN (https://cran.r-project.org/package=ruv). All computational source code required for reproducing all results presented in this paper are available in https://github.com/RMolania/NanostringNormalization.

## Data availability

The publicly available data sets used in this study are available through the following addresses: The TCGA lung cancer adenocarcinoma RNA-SeqV2 in the Broad GDAC Firehose (https://gdac.broadinstitute.org), data version 2016/01/28; Expression data for none-small cell lung cancer Affymetrix microarray data by Hou *et al.* [17], GEO GSE19188; Inflammatory bowel disease (IBD) study Nanostring data by Peloquin *et al.* [18], GEO GSE73094; Colon biopsies from UC patients and healthy controls Agilent microarray data by Noble *et al.* [21], GEO GSE11223; T cell study Nanostring data by Ye *et al.*[22], GEO GSE60341; T cell study Affymetrix microarray data by Ye *et al*. [22] GEO GSE60235; Dendritic study Nanostring data by Lee *et al*. [23], GEO GSE53165.

## Acknowledgements

T.P.S. was supported by Program Grant 1054618 from the National Health and Medical Research Council of Australia. A.D. was supported by funding from Cancer Australia for the lung cancer DNA repair study. R.M. was supported by two international Australian postgraduate scholarships (MIRS and MIFRS) from The University of Melbourne and a CTx PhD top up scholarship from the Cancer Therapeutics CRC. Dr Gavin Wright provided the lung cancer samples. We thank Dr Chun Jimmie Ye and Dr Gautam Goel for their comments on the near final draft. This study acknowledges support by the Victorian Government’s Operational Infrastructure Support Program to the Olivia Newton-John Cancer Research Institute. In particular, the authors thank groups who have made the raw and normalized data sets publically available.

## Competing interests

The authors declare no competing interests.

## Author contributions

R.M., A.D. and T.P.S. designed the overall approach. J. G. B and T. P. S developed the RUV-III method. R.M performed data analysis, A.D. provided data from the lung cancer study and carried out Nanostring lung study; R.M., J. G. B, A.D., and T.P.S. drafted the manuscript, which was revised and approved by all authors.

